# Alternative splicing plays a nonredundant role in greater amberjack (*Seriola dumerili*) in the adaptation to ambient salinity fluctuations

**DOI:** 10.1101/2024.01.03.574042

**Authors:** Chunyu Zhao, Yuqi Liu, Panpan Zhang, Xinhui Xia, Yuchen Yang

## Abstract

Alternative splicing (AS) is an important post-transcriptional mechanism for adaptation of fish to environmental stress. Here, we performed a genome-wide investigation to explore the biological importance of AS dynamics in greater amberjack (*Seriola dumerili*), an economical marine teleost species, in response to hypo- (10 ppt) and hyper-salinity (40 ppt) stresses. The results revealed high level of differential splicing in both gills and kidney upon the exposure to undesired salinity regimes. In gills, genes involved in energy metabolism, stimulus response and epithelial cell differentiation were differentially spliced in response to the deviation of normal water salinity, while sodium ion transport, erythrocyte homeostasis and cellular amide metabolism were enhanced in kidney to combat the adverse impacts of salinity changes. More importantly, the majority of the differentially spliced genes were not differentially expressed, and AS was found to regulate different biological processes from differential gene expression, indicative of the functionally nonredundant role of AS in modulating salinity acclimation in greater amberjack. Together, our study highlights the important contribution of post-transcriptional mechanisms to the adaptation of fish to ambient salinity fluctuations, and provides a theoretical guidance to the conservation of marine fishery resources under the increasingly extreme environmental challenges.

## 1 introduction

Occupying more than two-thirds of the Earth’s surface, the oceans are the largest ecosystem on the planet, hosting a huge number of organisms and playing crucial roles in maintaining the diversity and balance of life on Earth. However, in the recent few decades, global climate changes, such as global warming, glacier melting, and precipitation decreasing, have substantially changed the global water cycle and altered worldwide salinity distribution among different sea areas (Durack and Wijffels, 2010; Helm et al., 2011; Durack et al., 2012; Skliris et al., 2016; Ullah et al., 2021). One noticeable trend is that the freshwater is getting fresher while the saltwater becomes saltier (Durack et al., 2012). Till 2017, the mean salinity contrast between the low and high salinity regions had increased by more than 5% (Cheng et al., 2020). Furthermore, this imbalance has been found to be amplified by human activities (Omerspahic et al., 2022; Röthig et al., 2023). For instance, brine discharge during seawater desalination can lead to an abnormal increase in salinity of the receiving water body in coastal marine environment, while the enhanced urban drainage may freshen nearshore seawater (Omerspahic et al., 2022; Röthig et al., 2023). Salinity fluctuations in seawater would alter the dissolved oxygen level, nutrient concentration, and stratification of the oceans, thereby posting an adverse impact on the respiration, metabolism, reproduction, and even disease immunity of marine organisms (Lehtonen et al., 2016; Guo et al., 2020; Galkanda-Arachchige et al., 2021; Lu et al., 2022; Lacy et al., 2023; Röthig et al., 2023). For example, the increased water salinity was found to disrupt the normal energy allocation, and physiological and metabolic processes of fish, such as grass carp (*Ctenopharyngodon idella*) and African catfish (*Clarias gariepinus*), and finally inhibit their growth and survival (Smyth and Elliott, 2016; Zidan et al., 2022; Liu et al., 2023). Moreover, salinity change-induced biodiversity loss of plankton, macroalgae, coral reef, and seagrass may cause trophic collapse of marine ecosystem and lead to drops in fishery yield, which threaten ecosystem functions and food supply to human (Röthig et al., 2023).

*Seriola dumerili*, also known as greater amberjack, is a large saltwater bony fish of the family Carangidae. They are globally distributed in tropical and subtropical seas, including the Atlantic, Indian, Pacific, and Mediterranean Oceans (Zupa et al., 2017; Araki et al., 2018). Greater amberjack is an economically important fish that is widely popular for its tasty meat, and it is also a major predator and inhabitant of artificial reefs such as coral reefs and shipwrecks, maintaining the structure and function of these ecosystems (Pereira et al., 2023). It has been widely reported that the growth, development, and physiology of greater amberjack were sensitive to the changes of water salinity, temperature, and seawater pressure (Barany et al., 2021; Navarro-Guillen et al., 2022; Peng et al., 2022; Liu et al., 2023; Ru et al., 2023). Comparative transcriptomic analyses showed that salinity alternations substantially disrupt tissue growth and development, energy metabolism, and immune responses in greater amberjack (Peng et al., 2022; Liu et al., 2023). Correspondingly, muscle water content, gill and intestinal Na-K-ATPase activity, and growth-related processes were enhanced to combat the adverse impacts (Barany et al., 2021; Liu et al., 2023). However, most of the current researches only focused on the responsive regulations at transcriptional level. No attentions have ever been paid to the role of post-transcriptional mechanisms in modulating the resistance to seawater salinity fluctuations in greater amberjack.

Alternative splicing (AS) is one type of post-transcriptional regulatory mechanism in eukaryotes, and it can produce multiple isoforms with different combinations of exons, thus giving rise to multiple protein products from one single gene (Tian et al., 2022a). Therefore, AS can extend the diversity of structures and functions of proteins without increasing gene numbers. In fish, AS has been demonstrated to play critical roles in regulating the growth, development, and environmental adaptation of fish (Wan and Su, 2015; Tan et al., 2019; Long et al., 2020; Tian et al., 2020; Tian et al., 2022b; Xiao and Wang, 2022). The differential splicing of the genes related to spliceosome was indicated to assist the maintenance of gill homeostasis and improve the tolerance to hypoxia stress of spotted sea bass (*Lateolabrax maculatus*) (Ren et al., 2022). In another example, the expression of AS variants of *trim25* gene, which encodes an E3 ubiquitinated ligase of Tripartite Motif (TRIM) family, were up-regulated in Japanese flounder (*Paralichthys olivaceus*) in response to poly I:C treatment, indicative of their positively regulatory role of antiviral activity (Guo et al., 2022). Therefore, a comprehensive understanding of AS regulations associated with salt stress tolerance in greater amberjack allows us to fully understand the genetic basis underlying their physiological responses to fluctuating ambient salinity.

In this study, we performed a genome-wide investigation of AS dynamics in greater amberjack juveniles in response to the exposure to undesired salinity regimes. We further compared AS regulations to differential gene expression to explore their respective contributions to tolerance to water salinity fluctuations. Our findings will enrich our understanding of the molecular mechanisms underlying the adaptation to salinity stress in euryhaline fish, which provides theoretical guidance for the conservation of fishery resources under the increasingly extreme global climate change.

## 2 Materials and methods

### 2.1 Data preprocessing and sequence alignment

In our previous study, we sequenced the transcriptomes for two osmoregulatory tissues, gills and kidney, for greater amberjack juveniles reared under different salinity regimes: 10, 30 and 40 parts per thousand (ppt) (National Center for Biotechnology Information (NCBI) accession number: GSE220485) (Liu et al., 2023). Here, we re-used these RNA-seq data to investigate the role of AS regulations in greater amberjack in response to ambient salinity changes. Following the data preprocessing pipeline described in Liu et al. (2023), adapter contamination and low-quality reads were filtered out using Trim Galore v. 0.6.6 (Krueger, 2012) with a quality score cut-off of 25 and a read-length threshold of 50 bp. A genome index was constructed for the reference genome of greater amberjack (NCBI GCA_002260705.1) by the hisat2-built utility of HISAT2 package v. 2.2.1 (Kim et al., 2019). High-quality reads were mapped to the reference genome by the hisat2 program. For each sample, only the uniquely aligned reads were retained for downstream analysis. The alignment output was further assembled into full-length transcripts using StringTie v2.2.1(Kovaka et al., 2019). All sets of assembled transcripts were merged into a nonredundant set of transcripts using StringTie’s merge mode. The assembled transcripts were annotated by comparing against the reference annotations obtained from the NCBI RefSeq database using GFFcompare v. 0.11.2 (Pertea and Pertea, 2020).

### 2.2 Alternative splicing investigation and functional enrichment analysis

rMATS v. 4.0.2 (Shen et al., 2014) was employed to detect alternative splicing events of five major types: skipped exon (SE), alternative 5’ splice site (A5SS), alternative 3’ splice site (A3SS), mutually exclusive exons (MXE), and retained intron (RI) (Mehmood et al., 2020). For each event, the inclusion level of an alternative fragment in each sample was measured by “percent spliced in (PSI)”. The changes in average PSI values were computed between the hypo- (10 ppt) or hyper-salinity (40 ppt) treatment and the control group (30 ppt, CK) in gills and kidney, respectively, to investigate the shift in splicing patterns in response to ambient salinity alterations. For each comparison, the events with an FDR < 0.05 were considered to be significantly different between the two groups, and the genes with any differential splicing event were denoted as “DSGs”. Similar to other animals, SE events were the dominant AS type in greater amberjack, thus here we primarily focused on the functional importance of SE events. For the convenience, the genes with a larger PSI value of SE event under a treatment condition than CK were going to produce a higher proportion of the transcripts containing the alternative exon, and were denoted as “more exon-inclusive DSGs”, while those with a smaller PSI value were denoted as “less exon-inclusive DSGs”, which would give rise to more transcripts with the alternative exon skipped.

To access the functional importance of the DSGs, an OrgDb database was first built for greater amberjack using the R package AnnotationForge v. 1.38.1 (Carlson and Pages, 2017), based on the gene annotations obtained in our previous study (Liu et al., 2023). For both gills and kidney, Gene Ontology (GO) enrichment analysis was carried out for the DSGs via the enrichGO function of clusterProfiler package v. 4.6.2 (Wu et al., 2021). *P*-value < 0.05 was set as the criteria for statistical significance. For each tissue, the more exon-inclusive DSGs were compared across different salinity treatments, and the results were visualized by UpsetR package v. 1.4.0 (Conway et al., 2017). GO enrichment analysis was implemented for the DSGs 1) specific to 10 ppt saline treatment, 2) specific to 40 ppt saline treatment, and 3) identified under both treatments, respectively.

### 2.3 Premature termination codons (PTCs) induced by skipped exon events

One of the impacts of alternative splicing is the introduction of premature termination codons (PTCs) to transcribed isoforms, which may cause deficiencies of protein structure and function or even lead to nonfunctional proteins (Yeh and Hwang, 2020). PTCs account for approximately 20% of genetic defects in human diseases and are often associated with severe phenotypes (Oren et al., 2017). Here, we identified putative PTCs induced by SE events in greater amberjack and investigated their impacts on the product proteins. The coding sequence (CDS) regions were first predicted for each isoform based on the annotations of the assembled transcripts. Only those with unambiguous annotations of CDS, translation initiation codon, and termination codon were retained for downstream analysis. The coding sequences of each target isoform were extracted from the genome of greater amberjack, and all the putative termination codons were screened out in the coding regions. For the isoforms with multiple termination codons, the first appeared termination codon was considered as the PTC. The impact of PTCs on the structure of product proteins was assessed according to the distance between the PTC and the 5’ end of transcripts, where a small distance meant a large influence on the completeness of the protein structure.

### 2.4 Comparison between alternative splicing and differential gene expression

To further investigate the different roles between AS and gene expression in modulating salinity oscillation tolerance in greater amberjack, we compared the more exon-inclusive DSGs to the up-regulated differentially expressed genes (DEGs) under each treatment scenario. The DEGs were detected following the same criteria described in our previous study (Liu et al., 2023). GO enrichment analyses were then conducted for 1) the genes specifically regulated by differential AS (DS-specific genes), and 2) genes specifically regulated by differential expression (DE-specific genes).

## 3 Results

### 3.1 Differential splicing regulate salinity adaptation in gills of greater amberjack

In the gills of greater amberjack, compared to the CK group, RI events exhibited the largest alterations under both hypo- and hyper-salinity treatments, followed by SE and A3SS events (Figure 1B). When exposed to 10 ppt saline, 583 SE events were detected in 461 genes, where the transcriptional products of 182 genes displayed a higher inclusive level of the alternative exons, while the rest 279 genes were prone to skip the alternative exons. GO enrichment analysis of these more-exon inclusive genes revealed an enrichment of proton transmembrane transport and energy metabolic processes, such as glucose and alcohol metabolism, and sulfur compound biosynthesis (Figure 1C). Under 40 ppt salinity stress, 187 more exon-inclusive and 239 less exon-inclusive genes were identified compared to CK, where the more exon-inclusive genes were significantly overrepresented in the biological processes related to stimulus response, proton transmembrane transport, cilium assembly and epithelial cell differentiation (Figure 1D).

**Figure 1.**
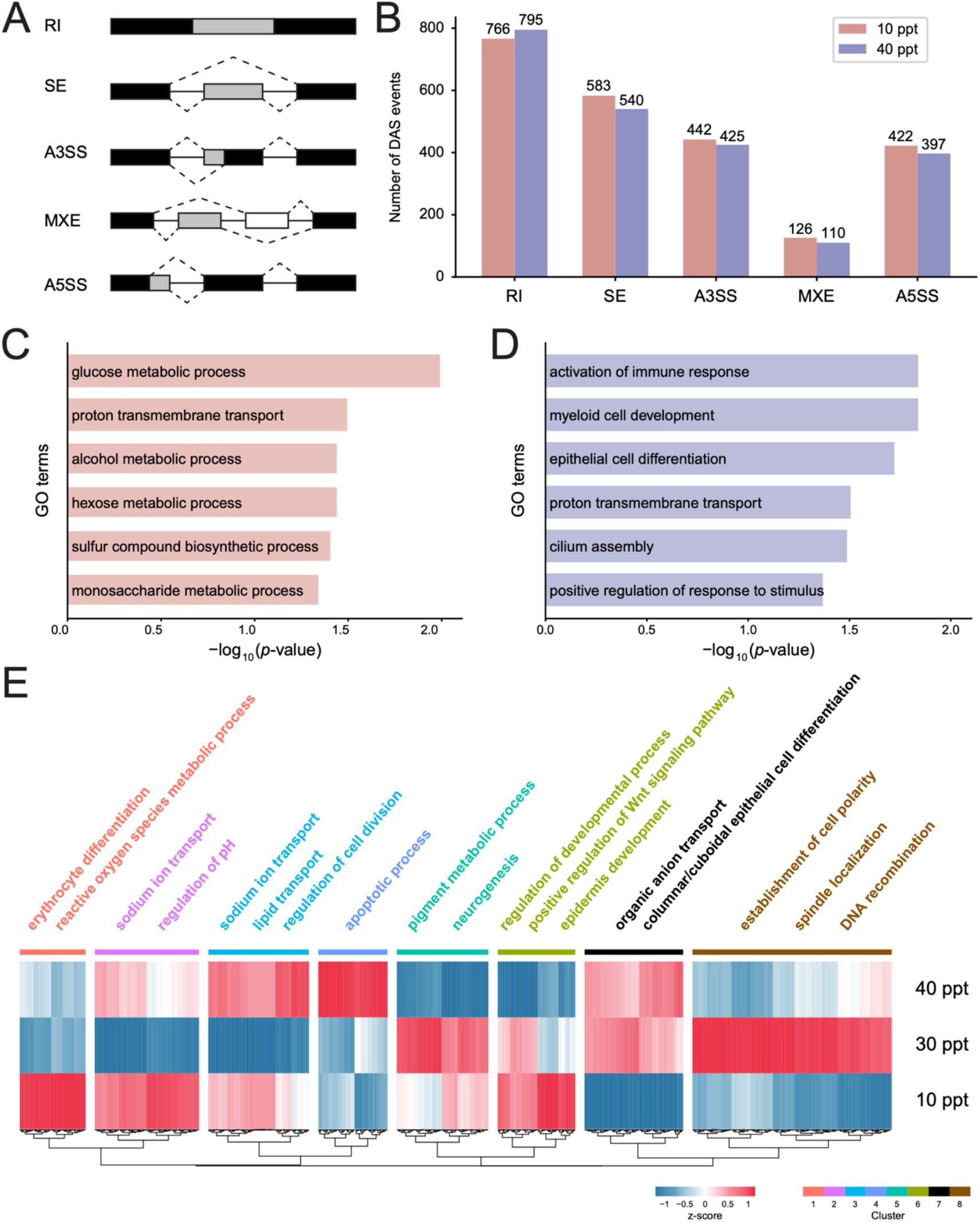
Alternative splicing reprogramming underlying salinity acclimation in gills of greater amberjack. (A) Schematic diagram showing five primary types of alternative splicing events we analyzed in this study. (B) Number of differential splicing events of each alternative splicing type in gills in response to hypo- (10 ppt) and hyper-salinity (40 ppt) stresses. (C) Representive Gene Ontology (GO) terms significantly enriched for the more exon-inclusive genes upon the treatment of 10 ppt saline. (D) Representive GO terms significantly enriched for the more exon-inclusive genes upon the treatment of 40 ppt saline. (E) Heatmap showing the percent spliced in (PSI) of the alternative exon of the exon-skipping genes across different salinity regimes. Representive GO terms enriched for the genes of each cluster are listed above the heatmap.

These DSGs were then grouped into 8 clusters according to their changes of PSI across the different salinity regimes (10, 30 and 40 ppt) (Figure 1E). In Cluster 1, 2 and 6, the majority of DSGs exhibited the highest inclusive level of the alternative exons in 10 ppt saline than the other two conditions. GO enrichment results showed that these genes were mainly involved in erythrocyte differentiation, reactive oxygen species (ROS) metabolism, sodium ion transport, pH regulation, regulation of developmental process, and positive regulation of Wnt signaling pathway. In contrast, the DSGs in Cluster 3 and 4 were of the highest exon inclusion in hyper-salinity treatment and these genes mainly participated in sodium ion transport, lipid transport, and regulation of cell division. These results suggested that greater amberjack activated sodium ion transport in gills to cope with the deviation of normal water salinity. A large group of DSGs displayed less exon inclusion under either hypo- or hyper-salinity than CK (Cluster 5 and 8), including those associated with pigment metabolic process, neurogenesis, cell polarity establishment, spindle localization, and DNA recombination.

### 3.2 Alternative splicing responds to salinity changes in kidney of greater amberjack

In the kidney, 759 significant SE events in 595 DSGs were detected under salinity stress at 10 ppt (Figure 2A). Of them, 276 DSGs displayed higher inclusion level of the alternative exons and these genes were found to be overrepresented in the GO terms of sodium ion transmembrane transport, organonitrogen compound catabolic process, regulation of pH, erythrocyte homeostasis, response to mechanical stimulus, myeloid cell homeostasis, and organophosphate metabolic process (Figure S1). While the 319 less exon-inclusive DSGs were highly representative in pigment granule transport, epithelial cell differentiation involved in kidney development, chromosome organization, and regulation of growth (Figure S1). Under 40 ppt salinity stress, 215 genes related to sulfur compound metabolism, aspartate family amino acid biosynthesis, lipid metabolism, and cellular amide metabolism process were found to include more exons than CK (Figure S2). Conversely, 287 genes were identified as less exon-inclusive DSGs and were enriched in regionalization, cell population proliferation, pigment metabolism, and epidermis development (Figure S2).

**Figure 2.**
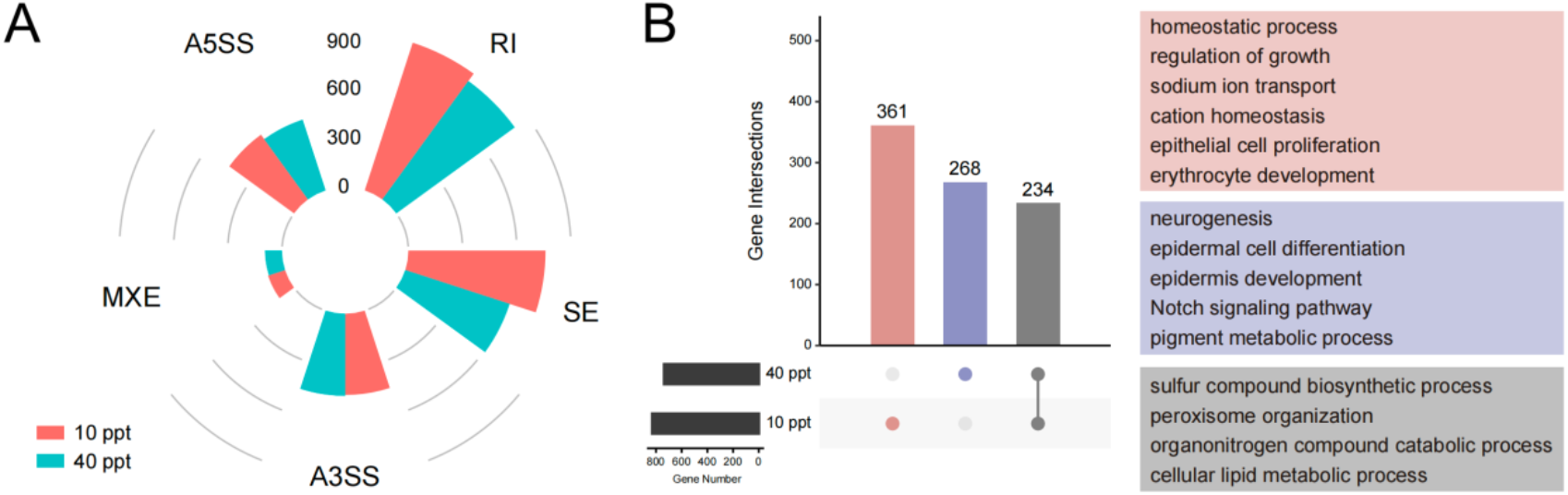
Differential splicing in the kidney of greater amberjack in response to salinity changes. (A) Number of differential splicing events of each alternative splicing type in kidney in response to hypo- (10 ppt) and hyper-salinity (40 ppt) stresses. (B) Overlap of the more exon-inclusive genes between 10 and 40 ppt saline treatments. The number of genes detected in each category was listed in the upset intersection diagram. Representive Gene Ontology (GO) terms significantly enriched for the genes of each category were listed at right, with the background color corresponding to the bars of the upset plot.

A cross comparison showed that, the exon inclusion level of 234 genes significantly increased under both 10 and 40 ppt saline treatments over CK, and these DSGs were mainly associated with sulfur compound biosynthetic process, peroxisome organization, organonitrogen compound catabolic process, and cellular lipid metabolic process (Figure 2B). Comparatively, 361 and 268 more-exon inclusive DSGs were specifically induced upon 10 and 40 ppt salinity stress, respectively (Figure 2B). Genes specifically responding to hypo-salinity stress were overrepresented in the GO terms of homeostatic process, regulation of growth, sodium ion transport, cation homeostasis, epithelial cell proliferation, and erythrocyte development, while those involved in hyper-salinity responses were enriched in epidermal cell differentiation, epidermis development, Notch signaling pathway, and pigment metabolism (Figure 2B).

### 3.3 Splicing-induced premature termination codons affect completeness of product proteins

Alternative cassette exons can moreover introduce multiple terminations to mRNAs and finally alter the coding potential of product proteins (Nikonova et al., 2020). In greater amberjack, SE events were found to give rise to PTCs under all four salinity treatments, which largely affect the completeness and function of the resulting proteins (Figure 3). It is worthy to note that, in gills, the majority of the introduced PTCs were located at the first 20% length of isoforms), that is, resulting in truncated protein products with a >80% loss in length (Figure 3). These transcripts were likely to be targeted by UPF factors and degraded by nonsense-mediated mRNA decay (NMD) pathway (Brogna and Wen, 2009). Comparatively, in kidney, a large proportion of PTCs occurred in the middle of the complete isoforms, indicating relatively smaller impacts on the functions of proteins.

**Figure 3.**
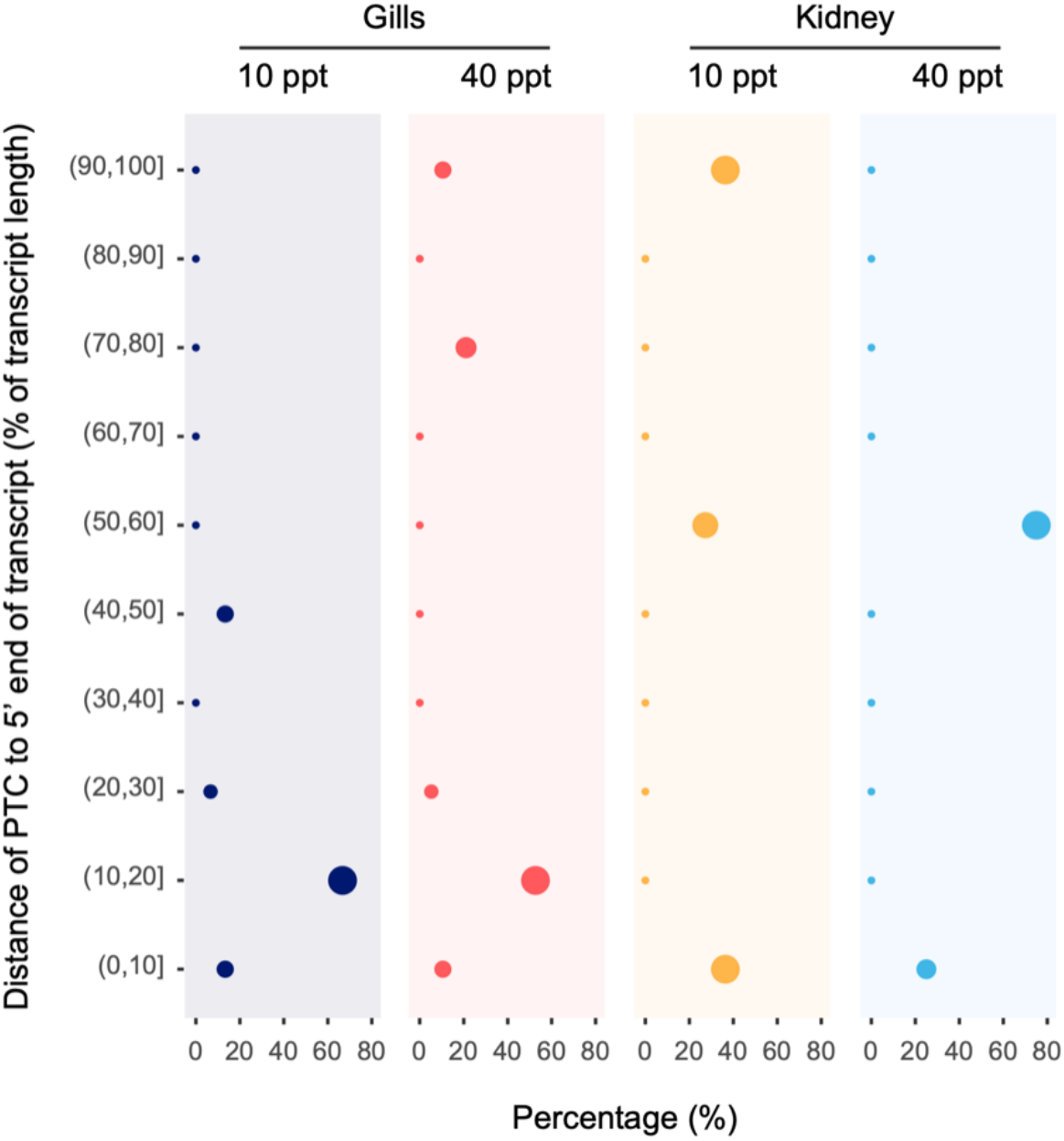
Distribution of PTCs in different locations on transcripts upon hypo- (10 ppt) and hyper-salinity (40 ppt) treatments in gills and kidney of greater amberjack. The location intervals of transcripts are listed along the vertical axis, and dot sizes correspond to the relative proportion of the PTCs in each location interval.

### 3.4 AS regulates different biological processes from gene expression under both hypo- and hyper-salinity

We then compared the differentially spliced genes to those of differential expression under each treatment to investigate their different roles in greater amberjack in mitigating the adverse impacts of salinity stress. For both gills and kidney, the majority of genes were either modulated by differential AS or differential expression, while only few genes were under the regulations of both mechanisms. More importantly, gene transcription and alternative splicing were found to participate in different physiological processes to enhance the tolerance of greater amberjack to water salinity oscillations. In the gills, when treated with 10 ppt saline, DS-specific genes were mainly involved in sulfur compound biosynthesis and proton transport, while the DE-specific genes were highly represented in the GO terms of growth, cartilage and osteoblast development, oxygen transport, and epithelium migration (Figure 4A). When exposed to 40 ppt salinity, the genes related to myeloid cell development, proton transport, and positive regulation of stimulus responses were mainly modulated by differential splicing, while those associated with cartilage development, extracellular matrix organization, regulation of BMP signaling pathway and response to external stimulus were differentially expressed (Figure 4B).

**Figure 4.**
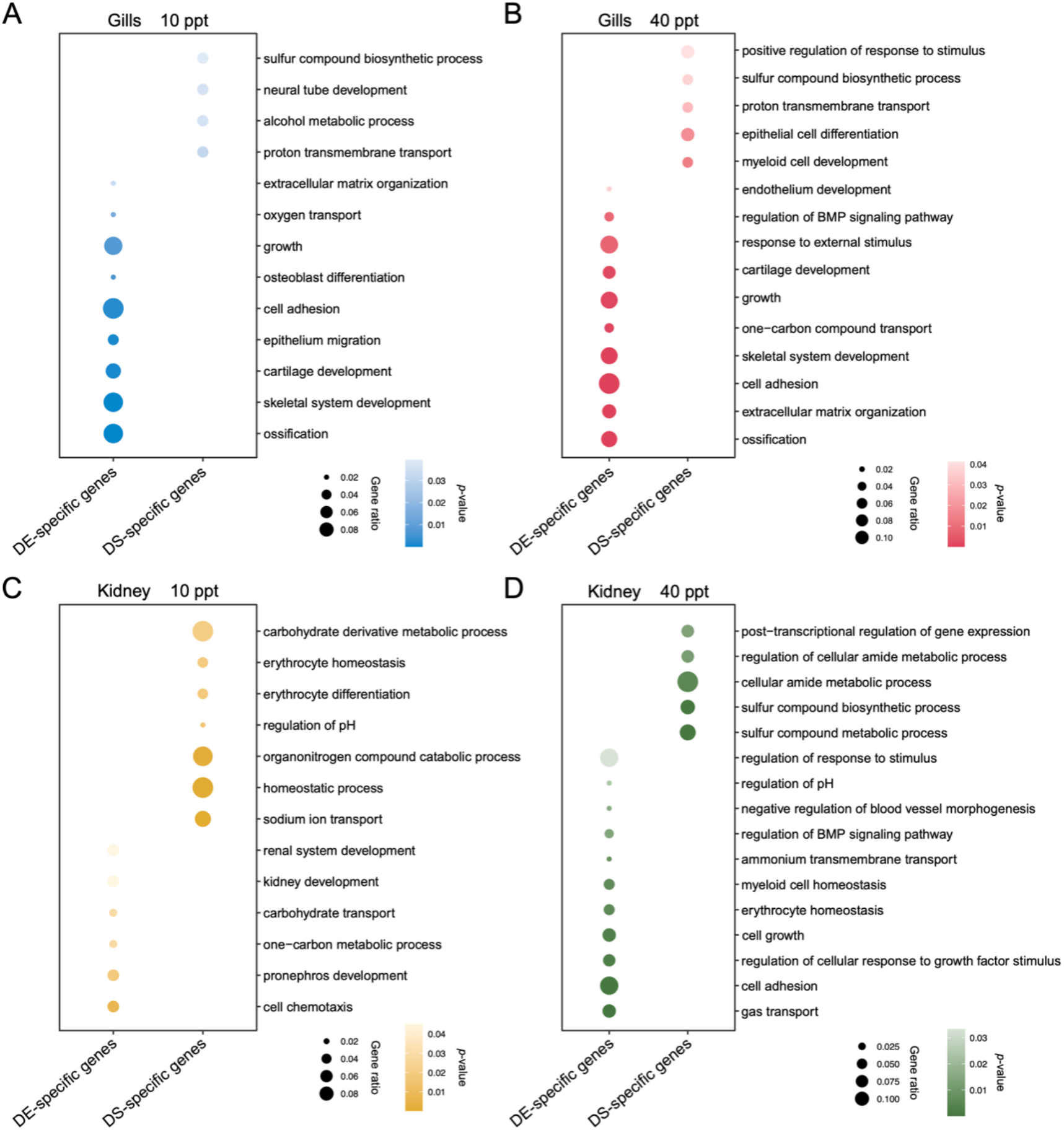
Representive Gene Ontology (GO) terms enriched for the genes that were specificially regulated by differential expression (DE-specific genes), and specifically regulated by alternative splicing (DS-specific genes).

In kidney, when exposed to low salinity environment, alternative splicing mainly affected the processes related to sodium ion transport, pH regulation, and erythrocyte homeostasis and differentiation, while transcription regulation was found to modulate the processes associated with kidney development (Figure 4C). Upon 40 ppt salinity, DSGs were significantly enriched in the GO terms of cellular amide metabolism, sulfur compound biosynthesis and metabolism, and post-transcriptional regulation of gene expression, while DEGs were mainly related to stimulus response, ammonium transport, pH regulation, and cell growth (Figure 4D).

## 4 Discussion

Water salinity is a key environmental factor affecting the yield of aquaculture (Callaway et al., 2012). In general, fish can only adapt to a limited range of water salinity changes; for instance, the optimum range of greater amberjack is 29 ‰−36 ‰ (Kültz, 2015; Ru et al., 2023). Deviation from the appropriate salinity range would disrupt their normal physiological and immune activities, which threatens their growth and survival (Zhang et al., 2017; Cui et al., 2019). Indeed, a previous study revealed a substantial up-regulation of the genes associated with organ development in both gills and kidney of greater amberjack juveniles to combat the adverse impact of salinity oscillations (Liu et al., 2023). In this study, we performed a genome-wide investigation of AS dynamics in greater amberjack juveniles in response to the exposure to undesired salinity regimes. The results revealed that both osmoregulatory organs, gills and kidney, were subjected to a high level of differential splicing during acclimation to hypo- and hyper-salinity stress, and these changes were mainly associated with sodium ion transport, energy metabolism, and cellular renewal and development, (Figures 1 and 2; Figure S1 and S2), which highlights the important role of AS in supporting the physiological reprogramming of greater amberjack to cope with short-term salinity fluctuation.

In gills, the genes involved in energy metabolism were significantly differentially spliced when exposed to 10 ppt saline (Figure 1C). For example, one of them encodes an enzyme hexose-6-phosphate dehydrogenase (H6PD) of endoplasmic reticulum. H6PD catalyzes the first two reactions of the oxidative branch of the pentose phosphate pathway: the conversion from glucose-6-phosphate and NADP^+^ to 6-phosphogluconate and NADPH (Senesi et al., 2010; Ge et al., 2020). The produced NADPH can reduce oxidized glutathione (GSSG) to reduced glutathione (GSH), which acts a key role in scavenging ROS and maintaining cellular homeostasis, especially upon environmental stresses (Senesi et al., 2010; Tomanek, 2012; Tomanek, 2015). Moreover, alcohol metabolism was also enhanced in the gills. For most saltwater fish, a decrease in water salinity alters their oxygen requirements in gills, thus may increase the production of lactate (Ern et al., 2014; Taugbøl et al., 2022). To combat the adverse impact of lactate buildup, crucian carp and goldfish (*Carassius*, Cyprinidae) can further covert lactic acid to ethanol and excrete it into ambient water via the special coupling of pyruvate dehydrogenase complex (PDC) and alcohol dehydrogenase (ADH) in gills, which largely improve their tolerance to low oxygen environment (Fagernes et al., 2017). However, no studies yet have investigated the detailed alcohol metabolic processes in greater amberjack, and future work is still required. When reared in a hyposalinity environment, the genes related to stimulus response, for example, *RNF111*, were activated in gills (Figure 1D). The coding product of *RNF111*, ring finger protein 111, was revealed to positively regulate DNA damage recognition in nucleotide excision repair, thus alleviating stress-induced DNA damage (Poulsen et al., 2013). Moreover, cilum assembly and epithelial cell differentiation were also differentially regulated by AS. Previous studies have revealed that cell proliferation and differentiation of gill epithelia were substantially activated in response to high salinity exposure, which can promote the osmoprotection of gill cells (Chretien, 1986; Fiol et al., 2009). Moreover, the leakiness of epithelial tight junctions was also increased to enhance the extrusion of sodium ions across the gill epithelium (Kültz, 2015). These findings highlighted the key role of epithelial remodeling in fish gills in hypo-osmoregulation.

In kidney, sodium ion transport and pH regulation were enhanced to combat the decrease in water salinity (Figure S1). Kidney is the main osmoregulatory organ in fish. The enhancement of ion transport in kidney can increase salt excretion and maintain the ion balance and blood osmolality in greater amberjack. Moreover, the splicing modes of the genes related to erythrocyte homeostasis were also significantly changed. A low ambient salinity may lower the internal osmolality of fish, which further alters the size and shape of red blood cells (Jensen et al., 2002). A rapid and efficient regulation of blood cell volume can guarantee the oxygen transfer between red blood cells and kidney. When exposed to hyper-salinity stress, cellular amide metabolic process, epidermis development, Notch signaling pathway, and pigment metabolism were activated to cope with the increased salinity (Figure 2B; Figure S2). The breakdown of amide/peptide bonds of proteins moreover gives rise to free amino acids, which are important cellular osmolytes in euryhaline fish (Aragão et al., 2010). A meta-analysis across 106 study cases showed that the cellular content of free amino acids was positively correlated with the elevation of salinity, highlighting their essential role in maintaining the homeostasis of osmotic pressure in fish in hyper-salinity environments (Huang et al., 2023). It is noteworthy that energy metabolism, including organonitrogen compound catabolism and lipid metabolism, were substantially activated by both 10 and 40 ppt saline treatments (Figure 2B). For both hypo- and hyper-salinity acclimation, extra energy costs are required by fish kidney for the osmoregulatory work, and the metabolism of lipids and proteins can serve as alternative fuel sources to enhance the energy supply (Aragão et al., 2010; Nguyen et al., 2016).

It is worthy to note that, in both gills and kidney, the most majority of differentially spliced genes were not differentially expressed, and these two mechanisms were found to modulate different biological processes (Figure 4). For example, in kidney, the genes involved in sodium ion transport and pH regulation were specifically differentially spliced upon 10 ppt salinity stimulus, while those associated with pronephros development were primarily under the regulation of differentially expression (Figure 4C). These indicated that AS is functionally nonredundant and plays a complementary role to gene transcription in regulating protein expression in greater amberjack in response to ambient salinity fluctuations. It is consistent with observations in other fish species, such as salmonid fish (*Salvelinus alpinus*) (Jacobs and Elmer, 2021)and spotted sea bass (Tian et al., 2020). AS and gene transcription are regulated mainly by independent mechanisms. The alternative procedures of pre-mRNA splicing are determined by the different recognitions and usages of splicing sites of spliceosome complex (Wang et al., 2015; Liu et al., 2022). In addition, the spliced mRNA products carrying PTCs would be targeted and degraded by the mRNA-surveillance mechanism NMD pathway (Stamm et al., 2005; Brogna and Wen, 2009). Our study also observed a large impact of PTCs on the structures and functions of the resulting proteins in both gills and kidney of greater amberjack (Figure 3). These features allow AS to extend and modulate the mRNA and protein repertoire of selectively constrained genes without altering their initial expressions, which acts as a key mechanism in the development of phenotypic plasticity and rapid ecological adaptation (Marden, 2008; Barbosa-Morais et al., 2012; Bradley et al., 2012; Jacobs and Elmer, 2021; Steward et al., 2022). Our current study adds further evidence for the nonnegligible role of AS in regulating stress tolerance in teleost fishes.

## 5 Conclusion

In this study, we provide a new insight into the role of alternative splicing in regulating physiological remodeling of greater amberjack to cope with the adverse impacts of fluctuated water salinity. The results show that alternative splicing plays a relatively independent role in fine-tuning protein expression in both gills and kidney of greater amberjacker in the early responses to salinity changes. Differential splicing primarily results in structural and functional changes of the proteins involved in sodium ion and proton transport, energy metabolism, and cell development and differentiation, which may improve the adaptation of greater amberjacker to salinity stress. These findings unravel the deeper side of the genetic basis underlying salinity acclimation in greater amberjack in response to the fluctuating marine environment, and provide a genomic view for the conservation of marine fishery resources.

## Acknowledgement

This research was financed by grants from the National Natural Science Foundation of China (32201420).

## Data availability statements

Data were obtained from the National Center for Biotechnology Information (NCBI) with accession number: GSE220485.

## Conflict of interst statement

The authors declare that the research was conducted in the absence of any commercial or financial relationships that could be construed as a potential conflict of interest.

