## Supplemental materials for "Alternative splicing plays a nonredundant role in greater amberjack (*Seriola dumerili*) in the adaptation to ambient salinity fluctuations"

* The authors contribute equally to this work.

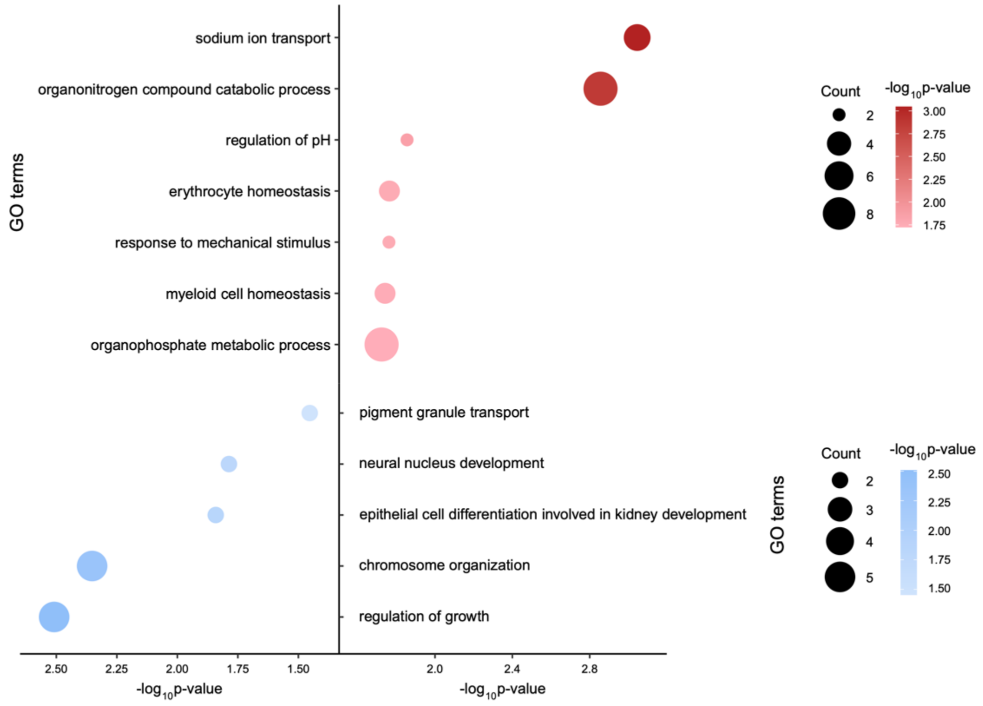


Figure S1. Representative Gene Ontology (GO) terms enriched for the more-exon inclusive (red) and less-exon inclusive genes (blue) in the kidney of greater amberjack in response to hypo-salinity (10 ppt) treatment.

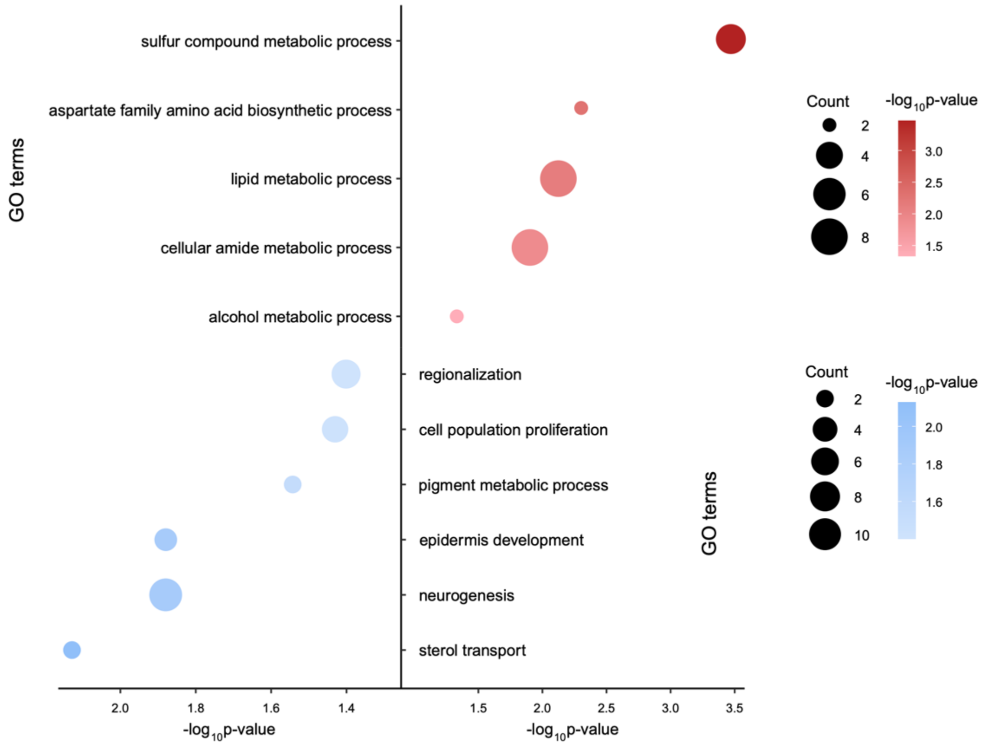


Figure S2. Representative Gene Ontology (GO) terms enriched for the more-exon inclusive (red) and less-exon inclusive genes (blue) in the kidney of greater amberjack in response to hyper-salinity (40 ppt) treatment.
